# Early and Late Components of EEG Delay Activity Correlate Differently with Scene Working Memory Performance

**DOI:** 10.1101/143461

**Authors:** Timothy M. Ellmore, Kenneth Ng, Chelsea P. Reichert

**Author notes:** Corresponding author (TME).

## Abstract

Sustained and elevated activity during the working memory delay period has long been considered the primary neural correlate for maintaining information over short time intervals. This idea has recently been reinterpreted in light of findings generated from multiple neural recording modalities and levels of analysis. To further investigate the sustained or transient nature of activity, the temporal-spectral evolution (TSE) of delay period activity was examined in humans with high density EEG during performance of a Sternberg working memory paradigm with a relatively long six second delay and with novel scenes as stimuli. Multiple analyses were conducted using different trial window durations and different baseline periods for TSE computation. Sensor level analyses revealed transient rather than sustained activity during delay periods. Specifically, the consistent finding among the analyses was that high amplitude activity encompassing the theta range was found early in the first three seconds of the delay period. These increases in activity early in the delay period correlated positively with subsequent ability to distinguish new from old probe scenes. Source level signal estimation implicated a right parietal region of transient early delay activity that correlated positively with working memory ability. This pattern of results adds to recent evidence that transient rather than sustained delay period activity supports visual working memory performance. The findings are discussed in relation to synchronous and desynchronous intra- and inter-regional neural transmission, and choosing an optimal baseline for expressing temporal-spectral delay activity change.

## Introduction

The ability to maintain information ‘online’ for a matter of seconds and then to use this information to realize immediate goals is critical for what is known as working memory [1]. While the behavioral limits of working memory, including capacity and duration, have been extensively documented [2, 3], it remains to be fully understood the neural dynamics that allow stimuli to be held in mind for short periods of time [3, 4].

Early work in non-human primates suggested [5] that it was the sustained and elevated firing of neurons in prefrontal and parietal cortices during the entire delay period since encoding but before a memory test that was important. More recent work challenges the long-standing view that elevated and sustained activity is necessary to support working memory [6]. Furthermore, work in monkeys suggests that working memory is supported by discrete and brief gamma and beta bursting [7]. A recent EEG study in humans showed that the active representation of an item in working memory can fall to baseline when attention shifts away. Afterwards a pulse of transcranial magnetic stimulation can cause the neural representation of the unattended item to reactivate as assessed using multivariate pattern analysis techniques [8]. This finding suggests that neural representations in working memory are dynamic and may be maintained by mechanisms that are ‘activity silent’ and do not necessarily involve sustained neuronal firing [9, 10].

Studies utilizing electroencephalography (EEG) in humans add to the complex picture regarding the sustained nature of oscillations during working memory delay periods [11–14]. Using scalp EEG, sustained synchronous alpha rhythms in parietal and occipital regions have been observed during the delay period of trials where the subject subsequently makes a successful probe choice [15, 16]. A more recent study points to beta activity, in addition to alpha activity, as playing a role in successful maintenance [4]. Oscillations in the theta range have been observed during the delay period as well [17]. The relationship of these electrical oscillations to widely used indirect measures of neural activity (e.g., BOLD-fMRI) remains a topic of ongoing investigation. Using intracranial EEG, which affords both high spatial and temporal resolution, delay period gamma band oscillation amplitude has been found to increase on electrodes located near fMRI activity, while theta band activity was elevated for electrodes located away from fMRI activity [18]. Synchronous frontal theta activity is reportedly also associated with active maintenance of information. Frontal theta activity is implicated in maintaining temporal information of encountered stimuli, whereas remembering item-specific details is associated with synchronous alpha and beta activity in parietal and occipital regions [19, 20].

A review of the literature on neural recording in non-human primates and on fMRI and EEG in humans shows that it remains to be completely characterized how neural activity evolves during the delay period of a working memory task. The long-standing elevated and sustained neural firing account is challenged by more recent studies that suggest transient dynamics [6, 21, 22]. Transient in this context means that activity could rise or fall at different times during the delay period. Activity could also be heterogeneous across frequency bands, with different bands exhibiting different relationships to memory performance. In the present study, the main questions addressed are: How does scalp EEG activity measured during the delay period of a working memory task change as both a function of time and frequency? Furthermore, how do the changes in delay activity predict subsequent short-term memory performance?

In the present study, participants completed several trials of a Sternberg task, a classic paradigm to assay working memory that when coupled with neural recording allows for a disambiguation of activity related to encoding, maintenance (during the delay period), and a probe of recognition memory for stimuli encountered before the delay. Here, the Sternberg paradigm was modified to use complex naturalistic scenes as stimuli. The use of complex scenes as visual stimuli served the dual purpose of having a large number of stimuli that are novel, and also to minimize the chance that participants could rely exclusively on a strategy of recruiting their phonological loop through covert articulatory rehearsal. The first and main objective was to analyze scalp EEG collected during the performance of the task in order to quantify delay period activity as a function of both time and frequency. The second objective was to understand how patterns of delay activity predict subsequent working memory performance by performing a correlation of the time-frequency delay period activity patterns with subjects’ ability to distinguish a probe stimulus presented immediately after the delay period.

To evaluate systematically the hypothesis of transient rather than sustained delay activity supporting working memory, a temporal-spectral evolution (TSE) analysis of EEG data was conducted with different baseline and temporal windows. TSE allows for quantification of event-related enhancement or suppression of EEG oscillation amplitude or power [23, 24]. In the present study, the event is the onset of the delay period and TSE is computed using different time windows relative to onset of the delay. Additionally, TSE is expressed as change relative to different baselines. For example, the entire delay period could serve as its own baseline and then each temporal unit of activity starting from the delay could be expressed relative to the mean activity taken across the delay. Alternatively, a period of time before or after the delay period could serve as the baseline, but then the resultant activity pattern would need to be interpreted relative to the sensory and cognitive processes occurring during each of these different baselines. The lack of systematic exploration in the literature motivates the final objective here of studying how changing temporal windows and baselines affects temporal-spectral measures of EEG delay activity, and the correlations between delay activity measures and memory performance.

## Materials and Methods

### Subjects

A total of 20 participants were recruited between October 13, 2015 and July 11, 2016 by flyers posted throughout the City College of New York campus. Included in the study were healthy adults between ages 18 and 42 with normal or corrected to normal vision and the ability to make button presses. Participants were excluded if they did not speak English or had a history of learning disabilities or sleep disorders. Each participant provided written informed consent and completed the study procedures according to a protocol approved by the Institutional Review Board of the City College of New York. Participants were compensated $15 per hour for participation. Five participants were excluded due to excessive noise in their EEG signals, leaving a total of 15 participants (11 males, age range 18 to 42, mean age 24.13 years, *SD* = 6.71) whose data are reported here.

### Task

Each participant completed a 100 trial modified Sternberg working memory task [25] with naturalistic scenes as stimuli. Scenes were 24-bit color images randomly sampled from the SUN database [26]. The SUN database contains tens of thousands of pictorial stimuli. Only a fraction of these (250) were sampled randomly from the SUN database and used in the present study. Care was taken to sample pictures of outdoor scenes with no clearly visible faces or people in the scenes. By using novel scenes, we sought to decrease chances of subjects using a default verbal rehearsal strategy, which would be easier to do with, for example, pictures of nameable objects. The task was programmed in SuperLab 5 (Cedrus Inc.). Each scene was displayed on a 27-inch LED monitor with a refresh rate of 60 hertz (Hz) and a screen resolution of 1920-by-1080. Participants sat 83.5 cm from the monitor and maintained stable viewing using a combined forehead/chin rest. Each scene measured 800-by-600 pixels on the screen, and from the subject’s point of view occupied a horizontal viewing angle of 17.2 degrees and a vertical viewing angle of 12.7 degrees. The experiment took place within a sound-attenuated booth (IAC acoustics) to minimize auditory and visual distractions.

Each trial of the working memory task consisted of the following phases (Fig 1): encoding of scenes, delay (maintenance), probe, and presentation of a phase-scrambled scene. During the encoding phase, two scenes were presented sequentially each for 2 seconds for a total of 4 seconds of encoding. During the delay (maintenance) period a white screen was visible to the participant for a total of 6 seconds. After the delay, a probe scene was presented for 2 seconds. If the scene was one of the two scenes in the immediately preceding encoding phase (50% chance) the participant was required to press the right (green) button on a RB-530 response pad (Cedrus Inc); if the probe scene was not one of the previous encoding set, the subject was required to press the left (red) button on the response pad. After the probe, a phase-scrambled image was presented for 5 seconds.

**Figure 1.**
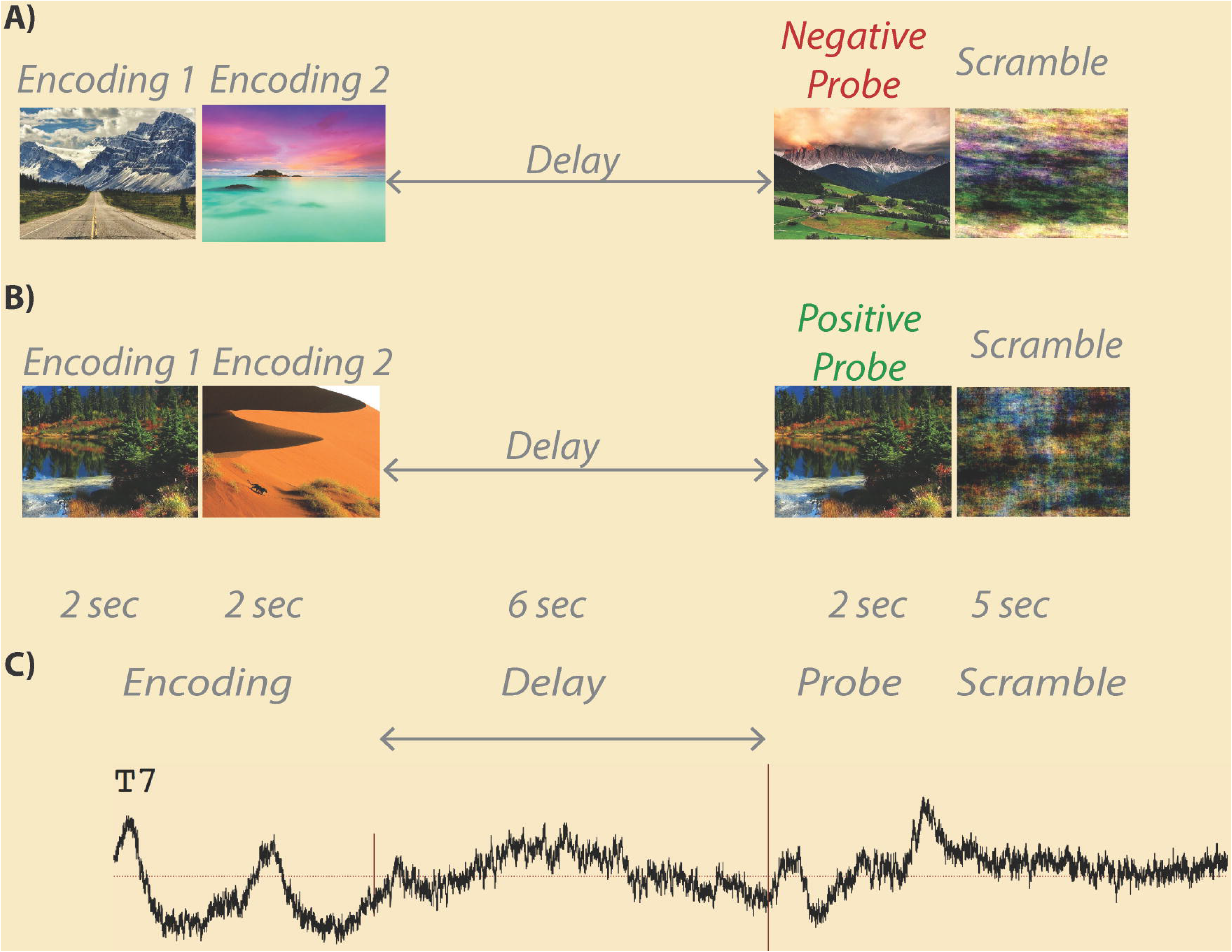
Example Scene Working Memory Trials. In a load 2 working memory trial, two encoding images are each presented for 2 seconds followed by a 6 seconds delay (maintenance) period. After the delay, a probe image is shown for 2 seconds followed by a 5 seconds scrambled scene baseline. Some trials contain a negative probe (a) where the probe stimulus is not one of the previously presented encoding images. Other trials contain a positive probe (b) where the probe stimulus is one of the previously presented encoding images. An example average EEG trace from one subject’s 94 artifact free trials is shown (c) with trial phases labeled. Onset of the delay period is marked by a short vertical red line, while offset of the delay period is marked by a longer vertical red line.

### Behavioral Analysis

Correct and incorrect trials were counted, and a signal detection analysis was performed to assess each participant’s performance. For the latter, a hit was counted when a scene from the immediately preceding encoding set was signaled by the subject by pressing a button indicating, correctly, that the stimulus had been previously seen (an old stimulus correctly classified as old). A false alarm was counted when a new scene not presented in the immediately preceding encoding set was indicated by the participant pressing a button indicating, incorrectly, that the scene had been previously presented (a new stimulus incorrectly classified as an old stimulus). For each participant, total hits and false alarms were expressed as proportions in each subject and used to compute a measure of sensitivity as the difference in standardized normal deviates of hits minus false alarms: d-prime (d’) = *Z(hit rate) – Z(false alarm rate)*. The d-prime sensitivity measure represents the separation between the means of the signal and noise distributions, compared against the standard deviation of the signal or noise distributions [27]. It was used as the primary covariate for the EEG analyses described below to determine how brain electrical activity during the delay period is related to a participant’s ability to distinguish stimuli presented during the last 10 seconds. The first 7 subjects were only presented 98 trials out of the planned 100 due to an error in task programming. Behavioral measures for these subjects were adjusted for this difference in trial number.

### EEG Acquisition

Electroencephalography (EEG) data were sampled at 1 kHz using Pycorder software from 62 scalp locations (Fig 2) using an active electrode system with an actiCHamp amplifier (Brain Products). Electrodes were placed at standard locations specified by an extended 10-20 system (https://osf.io/ebvsr/). The recording ground (Fpz) was located at the frontal midline and the recording reference was located at the left mastoid (TP9) leaving 61 scalp recordings (sensor level labels in Fig 3 and Fig 4). Two additional channels were designated for left (LOC) and right (ROC) vertical electrooculography (VEOG) recordings for subsequent isolation of eye blink artifacts. Recordings to disk were initiated after electrode impedances fell below 25 K Ohms. Although the standard convention is to reduce impedance to 5 K Ohms or below [28], the acquisition system utilizes active electrodes with noise reducing techniques built into the amplifier thereby ensuring that impedances under 25 K ohms are sufficient for interpretable signals [29]. Channels with impedance values above 25 K ohms were interpolated using data from neighboring electrodes with impedances below 25 K ohms. An auxiliary channel was used to record from a photosensor placed directly on a corner of the LED monitor. A 10-by-10 pixel square located under the photosensor was programmed to change from white to black during onset of each scene stimulus; it changed from black to white during stimulus offset. Recording the changing screen luminance from the photosensor signal at 1 kHz allowed for precise timing of stimulus onset and offset with respect to the recorded EEG data.

**Figure 2.**
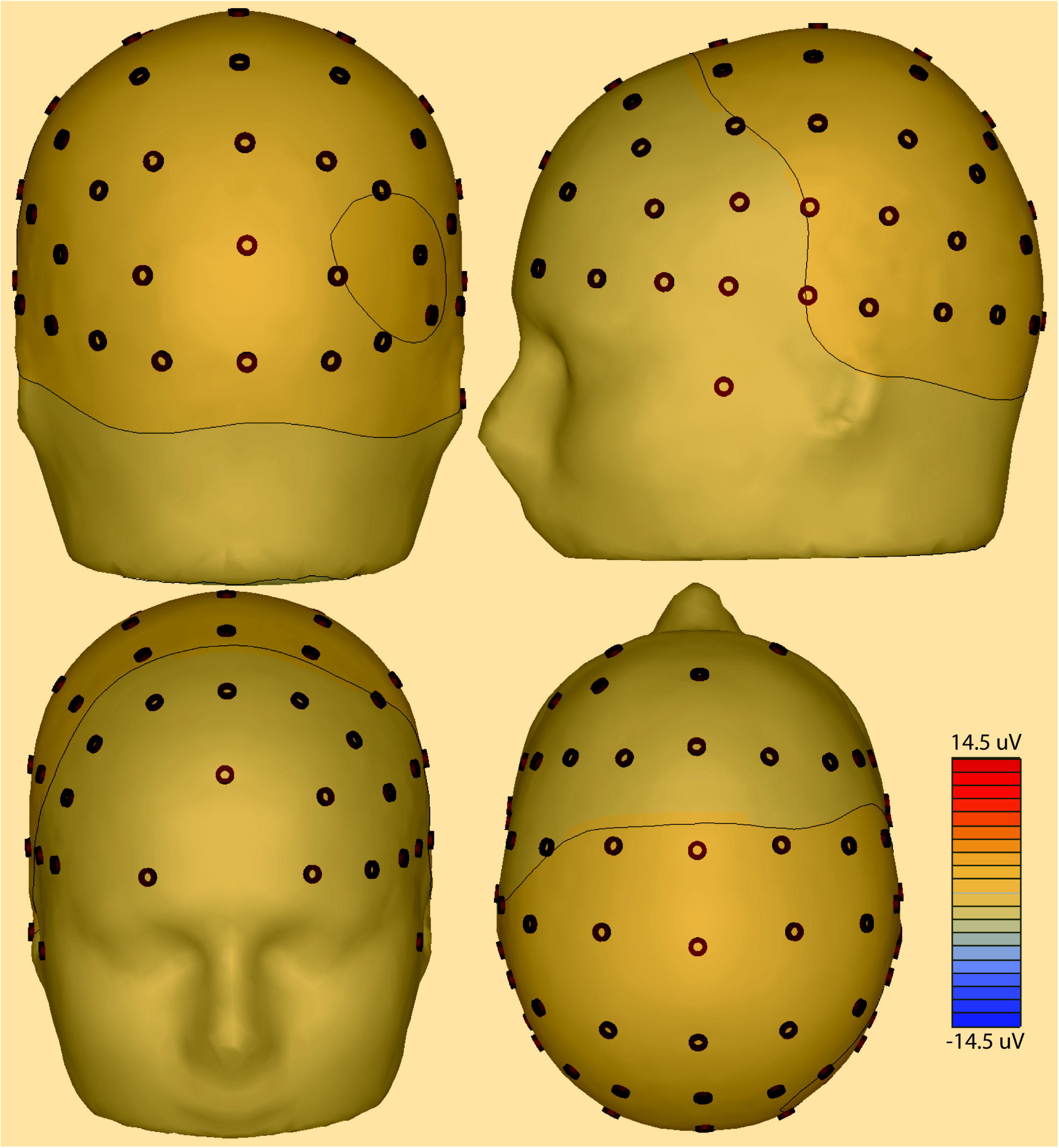
Scalp Electrode Montage Used for EEG Recordings. Scalp electrode positions used in the recordings are shown displayed on a head model. Contour lines represent 3D voltage amplitude mapping in a single subject 2000 ms after delay period onset. Note right posterior focus.

**Figure 3.**
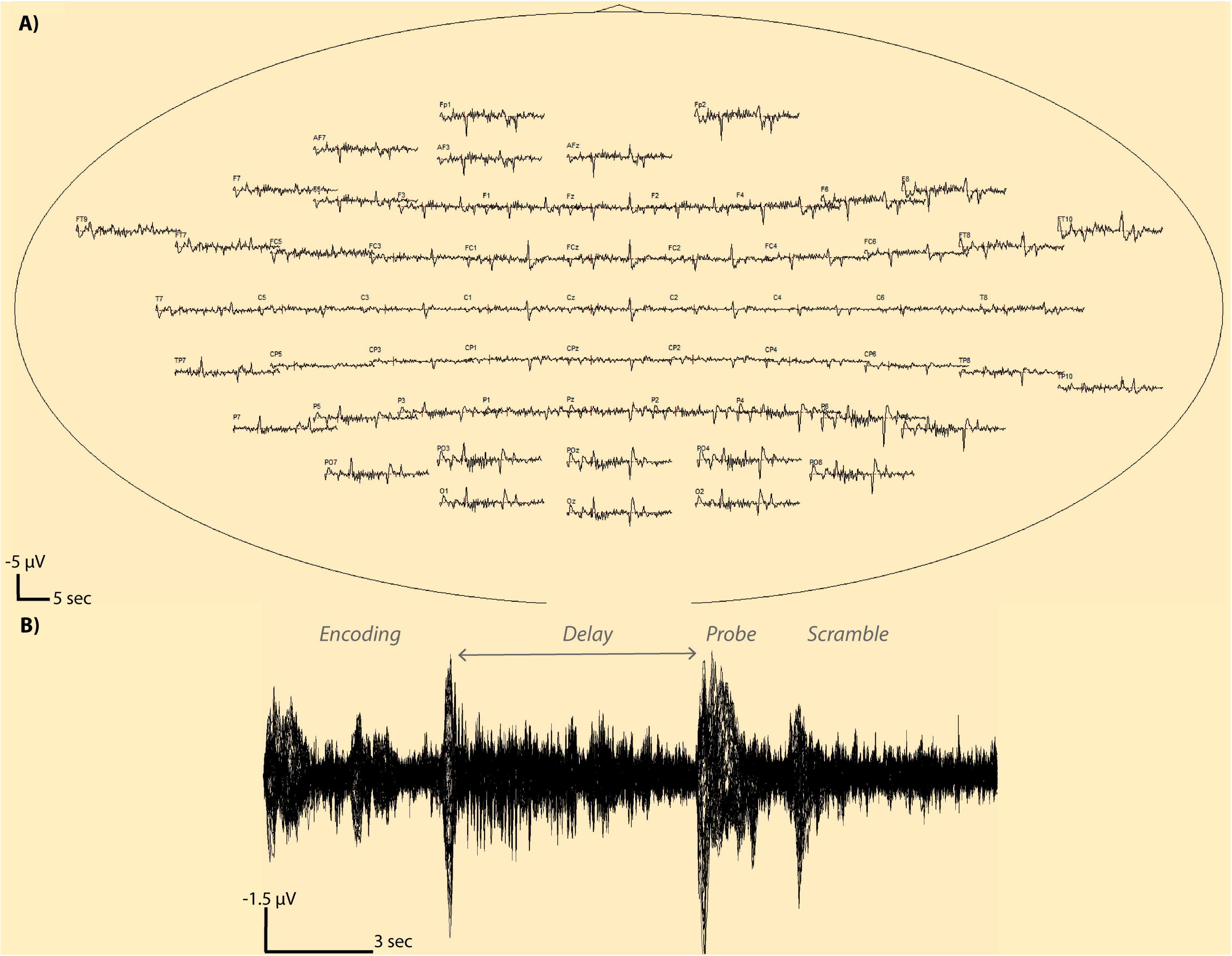
Averaged Whole Trial Window EEG Traces. An average EEG trace is shown at each sensor (a) for a single subject with 94 trials surviving artifact detection. Electrode T7 is shown in zoomed view (b) revealing the structure of the average EEG signal as a function of encoding, delay (maintenance), probe and scramble stimuli presentation. The T7 channel overplot view is also shown (c) showing superimposition of all 94 trial EEG traces from the beginning to end of the time window. Note that a negative potential is a upward deflection in these plots.

**Figure 4.**
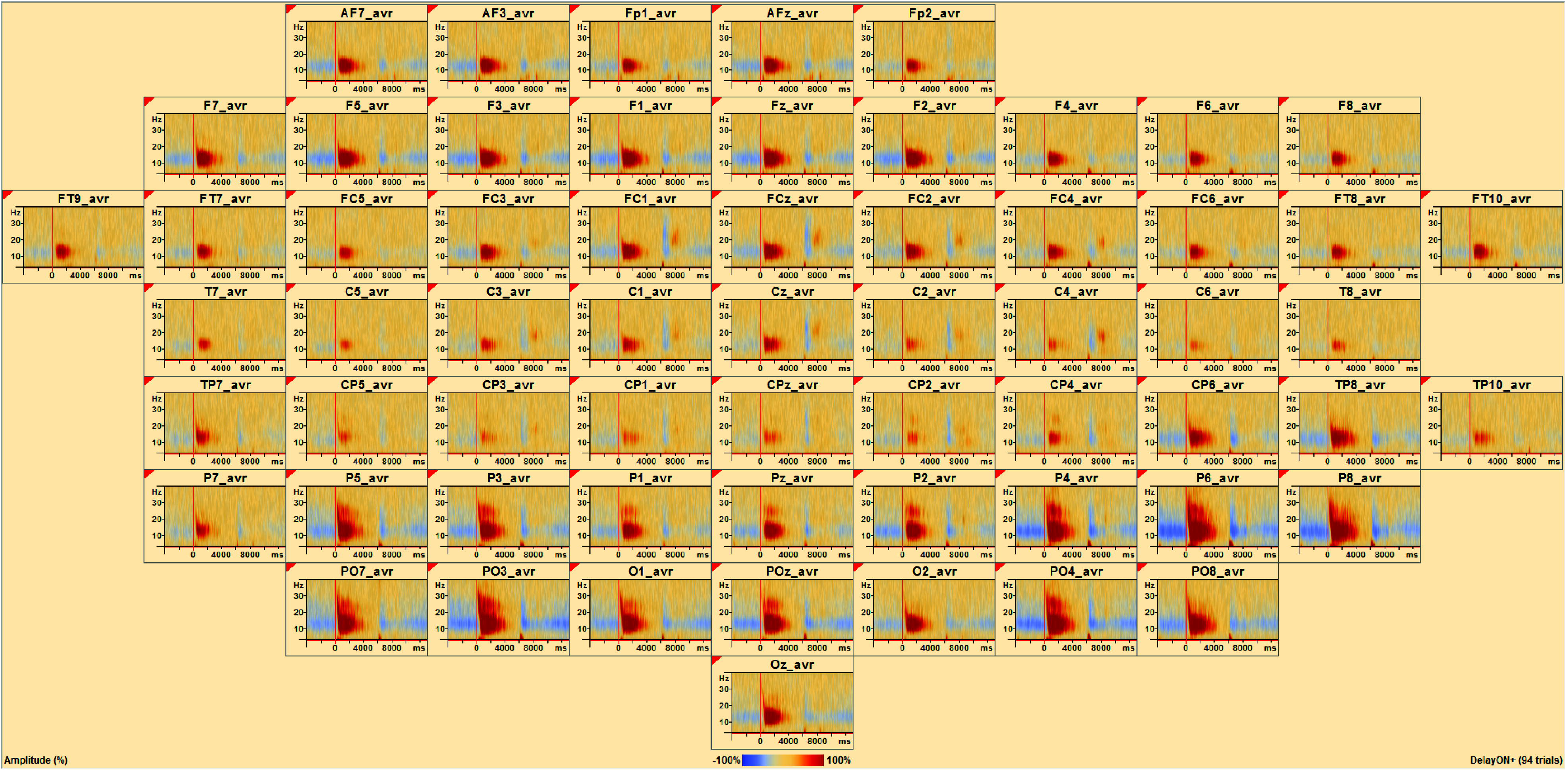
Temporal Spectral Evolution at the Sensor Level. At each sensor, the average TSE for a single subject with 94 trials surviving artifact detection is shown. Positive TSE change is visible after the onset of the delay period (vertical red line), while negative TSE change is evident before and after the onset and offset of the delay period. At each sensor, the x-axis of the TSE matrix shows time relative to the event, and the y-axis displays linearly scales with frequency. TSE amplitude intensities are color coded where blue is negative and red is positive. Note positive red change dominates during the delay period, whereas blue is most prominent during the encoding and probe phases.

### EEG Analysis

EEG signals were processed using BESA Research (v6.1) after re-referencing to a common average reference. First, notch (frequency 60 Hz, 2 Hz width) and bandpass (low cutoff 1 Hz, type forward, slope 6dB/oct; high cutoff 40 Hz, type zero phase, slope 12 dB/oct) filters were applied to all channels. Second, the signal on each channel was visually inspected to find, mark, and exclude the duration of all muscle artifacts. Third, a characteristic eye-blink was marked by finding an alternating deflection greater than 100 microvolts (*μ*V) between the LOC and ROC signals. Then a template matching algorithm was used to find all eye blink artifacts on all channels and remove the component of variance accounted for by the eye blinks [30, 31]. Finally, additional artifacts were isolated and excluded using amplitude (120 *μ*V), gradient (75 *μ*V), and low-signal (max. minus min) criteria (0.01). A participant’s data was used in further processing only if a minimum of 60% of trials survived this final artifact scan.

As detailed in [32], complex demodulation [33] was used as implemented in BESA to obtain the envelop amplitude and phase of each specific frequency band as a function of time. For each frequency of interest f_0_, the following steps were performed on single trial waveforms:

1. The original time-domain signal was multiplied by sin(2pf_0_t) and cos(2pf_0_t), respectively. This modulation operation shifts every signal at frequency f to the difference and sum frequencies (f±f0) in the frequency domain.
2. The two resulting signals were low-pass filtered to extract the frequency range that was originally centered around f_0_ and was shifted to the low frequency range f–f_0_. Thus, the low-pass cutoff frequency sets the half of the width of the frequency band for which the envelop amplitude and phase is computed. This filter implicitly defines the resolution in time and frequency. In this step, a finite impulse response (FIR) filter was selected with a low-pass frequency of 4 Hz (-6dB attenuation) corresponding to a full width at half maximum (FWHM) in the time domain of 130 ms.
3. The two output signals of the previous step define the real and imaginary part of a complex signal as a function of time. Its magnitude corresponds to half the envelop amplitude and its phase to the compound phase of the filtered frequency band centered at f_0_. In the following analyses and displays, we sampled f_0_ between 4 and 40 Hz in steps of 2 Hz and applied the FIR filter in sampling steps of 25 ms at latencies centered around the onset of the delay period.

After single trial waveforms were transformed into time-frequency space and the instantaneous envelop amplitude of brain activity was expressed as a function of frequency and latency, the change of amplitude with respect to the baseline was averaged over trials and displayed as a function of frequency and latency in temporal spectral evolution (TSE) event-related (de-)synchronization plots [23] using the following equation:

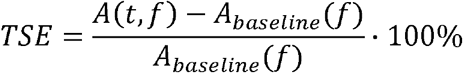

where *A(t,ƒ)* is the amplitude of activity during the specific timeframe of interest and frequency band and *Abaseline*(*ƒ*) is mean activity in a specific frequency band during a specified baseline period. The calculated value is expressed as a percent; the value is either a positive or negative change in spectral activity compared to the baseline period [34–36]. In other words, the amplitude for each time bin is normalized to the mean amplitude of the baseline epoch for that frequency. In the resulting matrix, the x-axis shows the time relative to the event, and the y-axis shows the frequencies. The TSE values in the matrix range from -100% to +infinity. A value of +100% means that activity is twice as high during the baseline epoch. Positive changes reflect event-related enhancement or synchronization as amplitude peaks around the event, while negative changes reflect event-related suppression or desynchronization as amplitude decreases around the event [35, 37].

In the present study, the timeframe (t) and baseline were systematically varied. In the first analysis, the timeframe was the entire 6 seconds delay period and the baseline was defined as the entire 6 seconds delay period. In the second analysis, the timeframe was the entire 17 second window starting at encoding through scrambled scene presentation with the baseline defined as the entire 17 second window. In the third analysis, the timeframe was the entire 17 second window starting at encoding through scrambled image presentation with the baseline defined as just the 4 seconds encoding phase of the task. In the fourth analysis, the timeframe was the entire 17 second window starting at encoding through scrambled scene presentation with the baseline defined as the 5 seconds scrambled scene presentation that came after the 2 seconds probe phase and before the next set of two scenes for the encoding period of the next trial. TSE calculations included all correct and incorrect trials surviving the eye-blink correction and artifact-scan, and for the sensor level analyses shown in Figs 6 and 7 the average TSE matrix across sensors is displayed.

The sensor level TSE analyses as described above were also carried out in source space [32, 38] using the BR_Brain Regions montage in BESA Research (Fig 5) and an age appropriate (20-24 year old) average adult head model [39]. The montage represents source space as 15 brain areas divided among left and right frontal, midline, parietal, temporal, and occipital regions.

**Figure 5.**
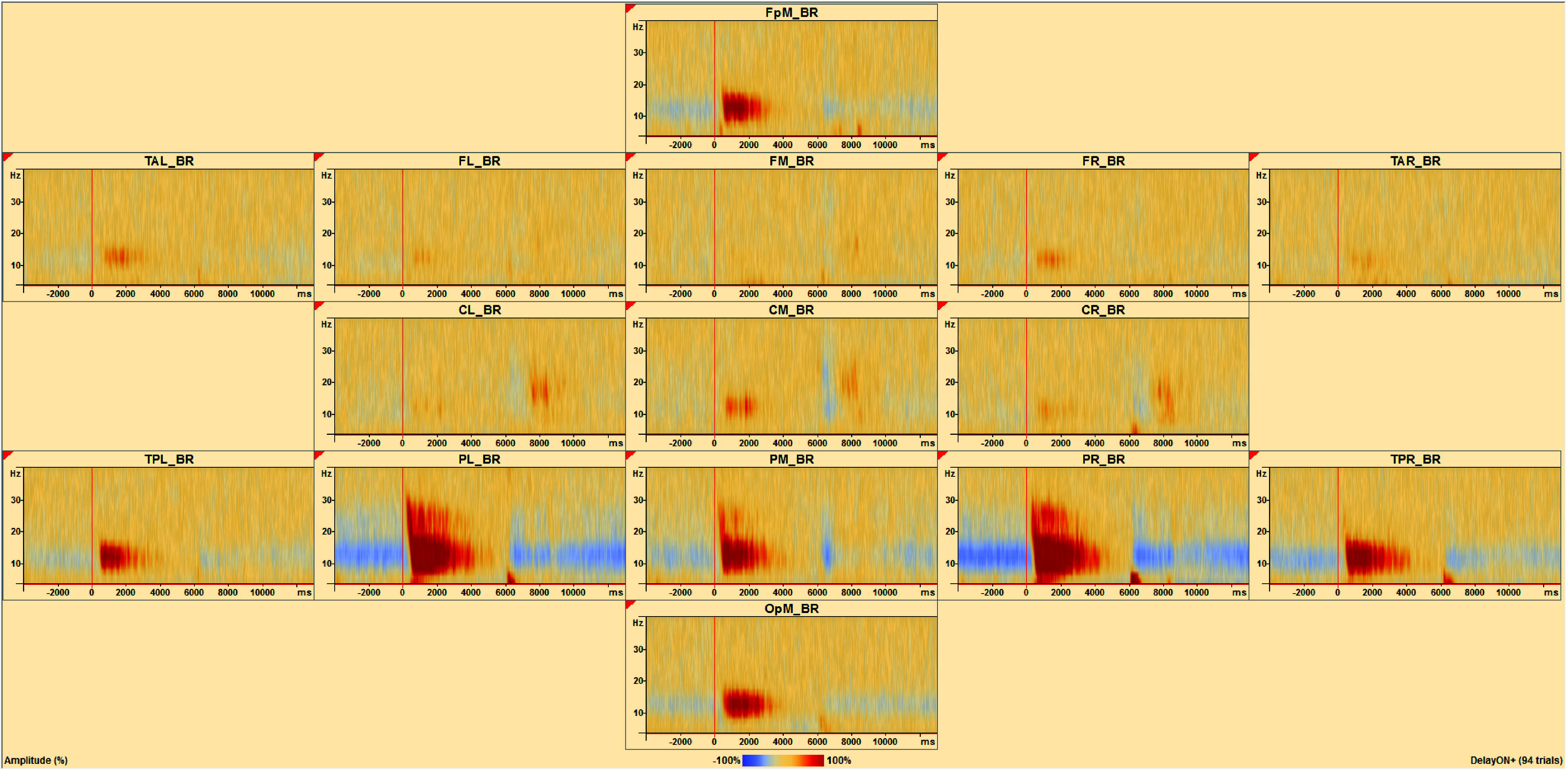
Temporal Spectral Evolution at the Source Level. A brain region source montage shows the average TSE for a single subject with 94 trials surviving artifact detection. Positive TSE change (red) is evident after the onset of the delay period (vertical red line), while negative TSE change (blue) is evident before and after the onset and offset of the delay period. These changes are particularly noticeable in medial frontal pole (FpM_BR), left posterior temporal lobe (TPL_BR), left parietal lobe (PL_BR), medial parietal lobe (PM_BR), right parietal lobe (PR_BR), right posterior temporal lobe (TPR_BR) and medial occipital pole (OpM_BR).

Statistical analyses of sensor and source space TSE matrices with participant d-prime performance measures was done in BESA Statistics v2.0 with corrections for multiple comparisons. Significant TSE-performance correlations are depicted in Figs 6 through 8 as dark red or blue shaded clusters overlaid on the average TSE activity matrices. Nonparametric cluster permutation tests (N=1,000) were computed in BESA Statistics v2.0 [40, 41]. For correlations, cluster permutation tests include two steps. First, spatially contiguous clusters of coherent *r* values exceeding a certain threshold along selected dimensions (time-by-frequency averaged over electrodes) are detected in the data. Here significant correlations were indicated by an *a priori* corrected p-value of less than or equal 0.05 (marginally significant by less than 0.10). Second, summed *r* values of the clusters are compared to a null distribution of *r* sums of random clusters obtained by permuting data order across subjects. This controls for type I errors due to multiple comparisons. Thus, the null hypothesis behind the permutation test for correlations assumes that the assignment of the covariate per subject (d-prime in this case) is random and exchangeable. Clusters of *r* values subjected to permutation are built across different dimensions. In the case of the sensor level analyses, clustering was performed on the time-frequency window after averaging over electrodes. In the case of the source estimation analyses, clustering was performed on 15 time-frequency windows each representing different source brain regions. The main idea behind data clustering done in combination with permutation testing is that if a statistical effect is found over an extended time period in neighboring channels, it is unlikely that this effect occurred by chance. A cluster value can be derived consisting of the sum of all r - or t-values of all data points in the cluster. For correlation r - or t-values, a statistical effect can have a positive or negative direction and thus positive and negative clusters can be found. The positive or negative cluster value is the test-statistic reported for each cluster, and the p-value reported is the one associated with that cluster based on permutation testing. For each of the 1000 permutations, new clusters are determined and the corresponding cluster values are derived for each cluster. Thus, a distribution of cluster-values can be established across all permutations. Based on this new distribution, the α-error of the initial cluster value can be directly determined. For example, if only 2% of all cluster values are larger than the initial cluster value, the initial cluster has a 2% chance that the null hypothesis was falsely rejected. This cluster would then be associated with a p-value of 0.02. Based on the computed cluster-value distribution, the probability of each initial cluster can be directly determined. The time and frequency ranges of each significant cluster were extracted to report locations with respect to well-known frequency bands, including theta (4-8 Hz), alpha (8-13 Hz), beta (13-40 Hz). The delta (<4 Hz) and gamma (40-90 Hz) ranges were not considered in this study.

## Results

### Behavioral

Participants performed well above chance on the task (percent correct *M* = 93.13, *SD* = 7.86; choice reaction time *M* = 1009 ms, *SD* = 91.66 ms). The mean number of hits (out of 50) was 45.87 (*SD* = 3.48), and the mean number of false alarms was 0.87 (*SD* = 1.77). The resulting average sensitivity, d-prime, to distinguish old from new scenes was 3.70 (*SD* = 0.70).

### Sensor Level TSE

### Delay Window Baseline

The average TSE matrix including artifact-free correct and incorrect trial data averaged over sensors is shown for the delay period window (Fig 6a). An area of negative change, relative to the whole delay window baseline, can be seen spanning 6 to 28 Hz, including from the start of the delay period, delay onset time=0, to 600 ms. A weak positive change from baseline is present from around 500 ms to 2925 ms spanning frequencies 6 to 26 Hz. A stronger positive change is present in the 4 to 8 Hz range from around 275 ms to 1650 ms. Finally, widespread, weak negative change is present from approximately 4125 ms to the end of the maintenance period spanning the 8 to 26 Hz range and also from 2500 ms to 5475 ms spanning the 4-8 Hz range. Statistical analysis of the correlation between TSE and participant d-prime measures for this window and baseline revealed three significant clusters (Fig 6b). The most significant cluster, Cluster 1, occurred from 4775-6000 ms, spanned a frequency range of 4 to 36 Hz, and contained activity that was negatively correlated (Fig 6e, *Cluster Value = -993.061, p* = .017) with d-prime. The next most significant cluster, Cluster 2, occurred from 1400 to 2450 ms, spanned a frequency range of 4 to 24 Hz, and was positively correlated (*Cluster Value = 665.685, p* = .022) with d-prime. The third most significant cluster, Cluster 3, occurred from 3850 to 4800 ms, spanned a frequency range of 8 to 26 Hz, and was negatively correlated (*Cluster Value = - 640.877, p* = .032) with d-prime.

**Figure 6.**
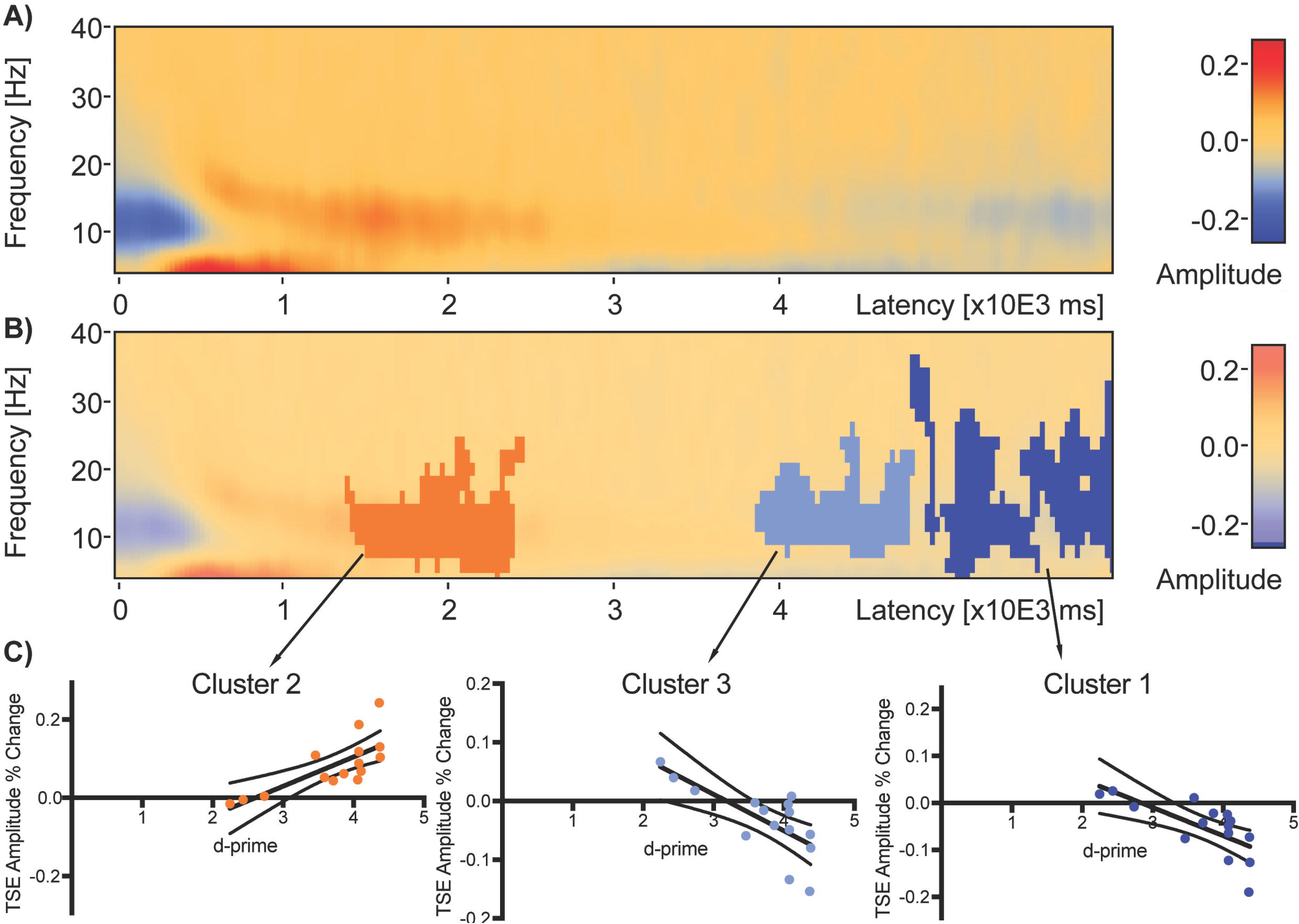
Average Delay Window TSE and Significant Clusters of Correlation with Performance Across All Subjects. Average delay period TSE plot (computed using whole delay window as baseline) with no statistical significance overlay (a) shows initial negative change from 6 to 28 Hz at the start of the delay period 0 to 600 ms. Weak positive change (red) is then present from around 500ms to 2925 ms at 6-26 Hz. Stronger positive change is present in the 4-8 Hz from around 275 ms to 1650 ms. A switch to negative change (blue) is present from approximately 4125 ms to the end of the delay period in the 8-26 Hz range and from 2500 ms to 5475 ms in the 4-8 Hz range. Correlation of TSE with subject d-prime revealed three significant clusters (b) in descending order from most significant. Significant clusters are represented by darker shades of red or blue overlaid on TSE activity. Cluster 1 (c, latency 4775-6000 ms and frequency range 4-36 Hz) and Cluster 3 (d, latency 3850-4800 ms and frequency range 4-24 Hz) were negatively correlated with d-prime (Cluster 1: *Cluster Value = -993.061*, p = .017; Cluster 3: *Cluster Value = -640.877, p* = .032) and Cluster 2 (e, latency 1400-2450 ms and frequency range 4-24 Hz) was positively correlated with d-prime (*Cluster Value = 665.685, p* = .022). Dark black lines on the graphs (c-e) are linear regression fits and dotted lines are 95% confidence intervals.

### Whole Window Baseline

The average TSE matrix including artifact-free correct and incorrect trial data and averaged over sensors is shown for the entire trial window, from the start of encoding to the end of the scrambled image presentation (Fig 7a). The baseline for this analysis is the entire trial window. The delay period is indicated by 0 to 6000 ms on the x-axis. A period of positive change is noticeable from approximately 425 ms to about 5050 ms ranging from 6 to 26 Hz. There is a short period of weaker positive change between 275 ms to 1950 ms ranging from 4 to 7 Hz. Weak negative change occurs between 0 and 400 ms ranging from 4 to 20 Hz and also from 5150 ms to the end of the delay period at 6000 ms in the 4 to 7 Hz range. Statistical analysis of the correlation between TSE and participant d-prime measures revealed one significant cluster ranging from 1400 to 3150 after delay onset, spanning frequencies 4 to 28 Hz, and that was positively correlated with d-prime (*Cluster Value = 1109.9, p* = .041).

**Figure 7.**
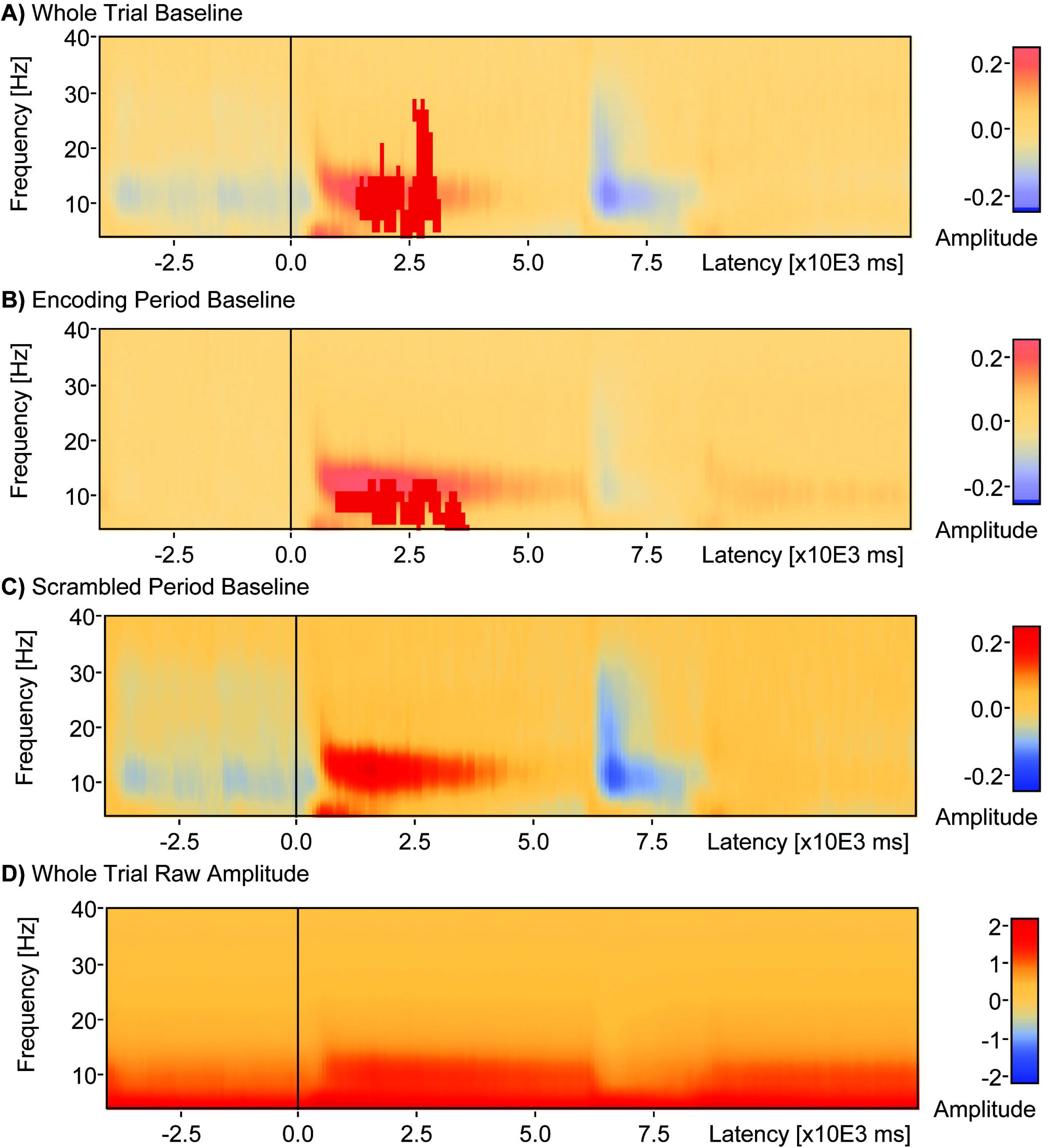
Effect of Baseline Window on TSE-Performance Correlations. When the whole 17 seconds window from encoding to scrambled scene presentation was used as baseline in computation of TSE (a) a single cluster (latency 1400-3150 ms and frequency range 4-28 Hz) was found that positively correlated with d-prime (*Cluster Value = 1109.9, p* = .041). When the 4 seconds period of encoding was used as baseline in computation of TSE (b) a single cluster (latency 900-3700 ms, frequency range 4-12 Hz) was found that correlated positively with d-prime (*Cluster Value = 830.1, p* = .024). When the 5 seconds period of scrambled scene presentation after the probe stimulus was used as the baseline in computation of TSE (c), there were no significant clusters found that correlated with d-prime.

### Encoding Period Baseline

The average TSE plot including artifact-free correct and incorrect trial data averaged over sensors is shown for the entire trial window, but with only the encoding period serving as baseline (Fig 7b). A positive change is visible after the onset of the delay period from 450 ms to about 6000 ms spanning the 8 to 26 Hz range, with a short period of weaker positive change between 325 and 1875 ms spanning the 4 to 8 Hz range. Statistical analysis of the correlation between TSE and participant d-prime measures revealed one significant cluster (Fig 7b) ranging from 900 to 3700 ms after delay onset, spanning frequencies 4 to 12 Hz, and positively correlated with d-prime (*Cluster Value = 830.1, p* = .024).

### Scrambled Scene Baseline

The average TSE matrix including artifact-free correct and incorrect trial data averaged over sensors is shown for the entire trial window, but with only the post-probe scrambled scene period serving as baseline (Fig 7c). There is a visible positive change from 375 ms to about 5250 ms in the 8 to 26 Hz range and from 275 ms to about 2000 ms in the 4 to 7 Hz range after the onset of the delay period. There is weak negative change from 0 to 425 ms in the 4 to 22 Hz range, and also from 5150 ms to the end of the delay period in the 4 to 7 Hz range. Statistical analysis of the correlation between TSE and participant d-prime measures revealed no significant clusters.

### Source Level TSE

The average TSE matrix including artifact-free correct and incorrect trial data is shown for the entire trial window, from the start of encoding to the end of the scrambled scene presentation, for source signals representing 15 brain regions (Fig 8d). The baseline for this analysis is the entire trial window. Strong positive change was evident after the onset of the delay period, while negative change was evident during the encoding period (before the delay) as well as during the probe period after the delay. This pattern of changes was particularly noticeable in the medial frontal pole, left and right posterior temporal lobe, the parietal lobe, and the medial occipital pole. Statistical analysis of the correlation between the 15 source TSE signals and participant d-prime measures revealed one cluster (Fig 8b, blue) ranging from -3750 to +275 ms, spanning 4 to 40 Hz, that was significantly negatively correlated with d-prime (*Cluster Value = -3223.92, p* = .018). A second cluster (Fig 8b, red) was also revealed, ranging from 1225 to 3150 ms and spanning 4 to 26 Hz, that was positively correlated (*Cluster Value = 1097.58, p* = .06) with d-prime and reached marginal significance. These clusters were located in the right parietal source region (PR_BR, region of source signal depicted orthogonally in Fig 8a). The cluster of negative correlation occurred during the encoding period of the task, while the cluster of positive correlation occurred during the delay period of the task. An additional source analysis was done with a restricted window spanning just the delay period, with the same window of time serving as baseline. Statistical analysis of the correlation between the 15 source TSE signals and participant d-prime measures revealed one cluster (Fig 8d) ranging from 1400 to 2450 ms, spanning from 4 to 26 Hz, and that was significantly positively correlated with d-prime (*Cluster Value = 680.062, p* = .021) after a multiple comparison correction including the number of source regions. This cluster was also located in the right parietal source region.

**Figure 8.**
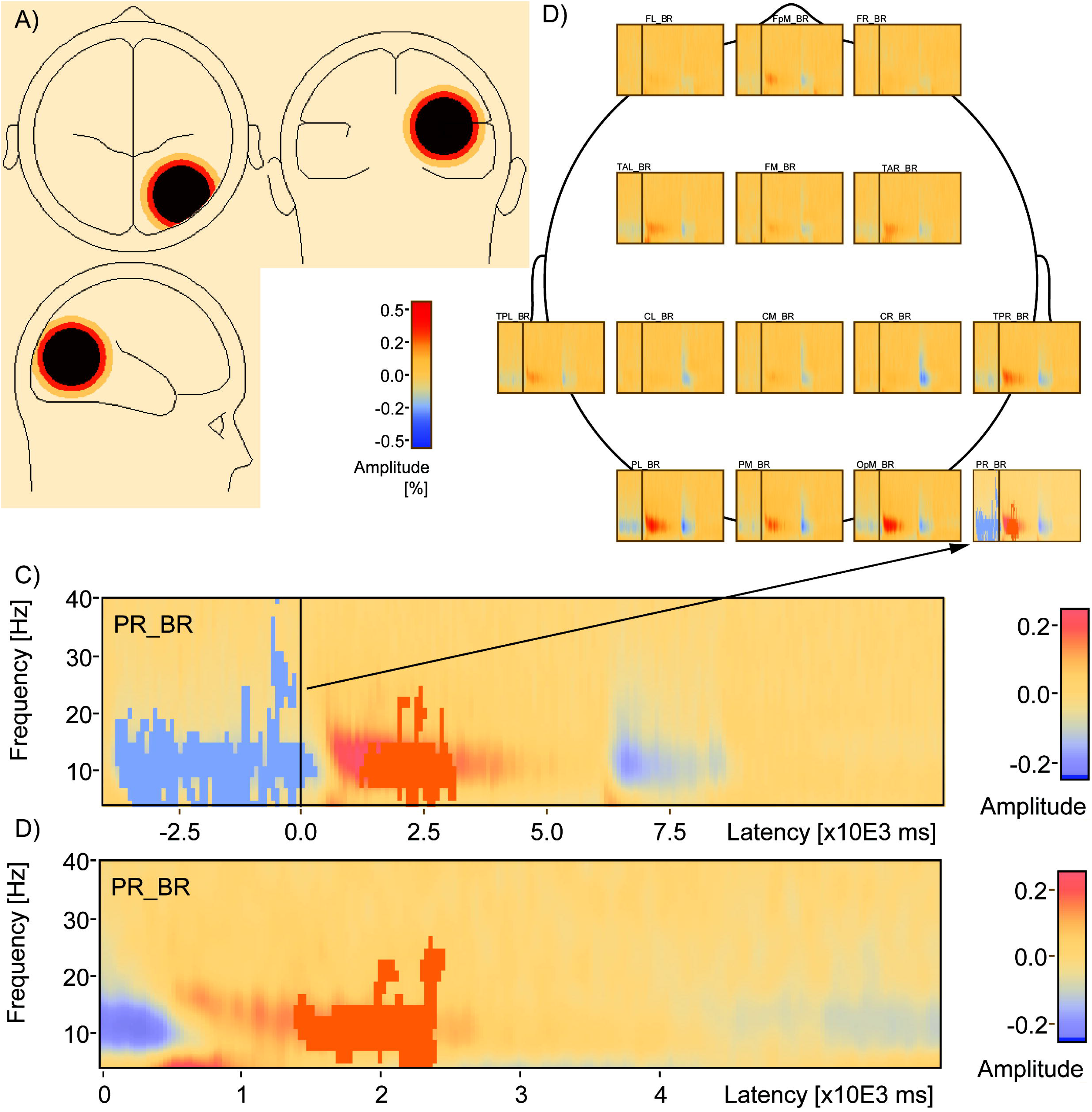
Source Space Analysis Reveals TSE-Performance Correlation in a Right Parietal Region. Source TSE analysis finds that PR_BR (a) includes a cluster of negative correlation with d-prime (blue, *Cluster Value = -3223.92, p* = .018) during encoding and a positive correlation with d-prime during the delay (red, *Cluster Value = 10.97.58, p* = .06) in a whole window analysis (b) and a cluster of positive correlation with d-prime (red, *Cluster Value = 680.062, p* = .021) in a smaller temporal window that included just the delay period (c). Location of the PR_BR relative to the other sources used to compute TSE is shown in a top down view (d).

### Amplitude Variance

One hypothesis for why there are fewer clusters of significant correlations with performance when the analysis window is expanded (Fig 7 versus Fig 6) is that amplitude variance increases as window length increases thereby reducing statistical power. To explore this hypothesis, the log10 raw amplitude variance was computed for each subject within two windows: 1) the whole 17 sec trial window and 2) the 6 sec delay period. Variance was found to decrease as a function of frequency, but was greater in magnitude across frequencies for the whole trial window compared to the delay period window (Fig 9a). Subject differences in log10 raw amplitude variance (whole trial window minus delay period window) were greater than zero in a frequency range between 4 and 40 Hz, with local peak differences at 8 Hz and 22 Hz (Fig 9b).

**Figure 9.**
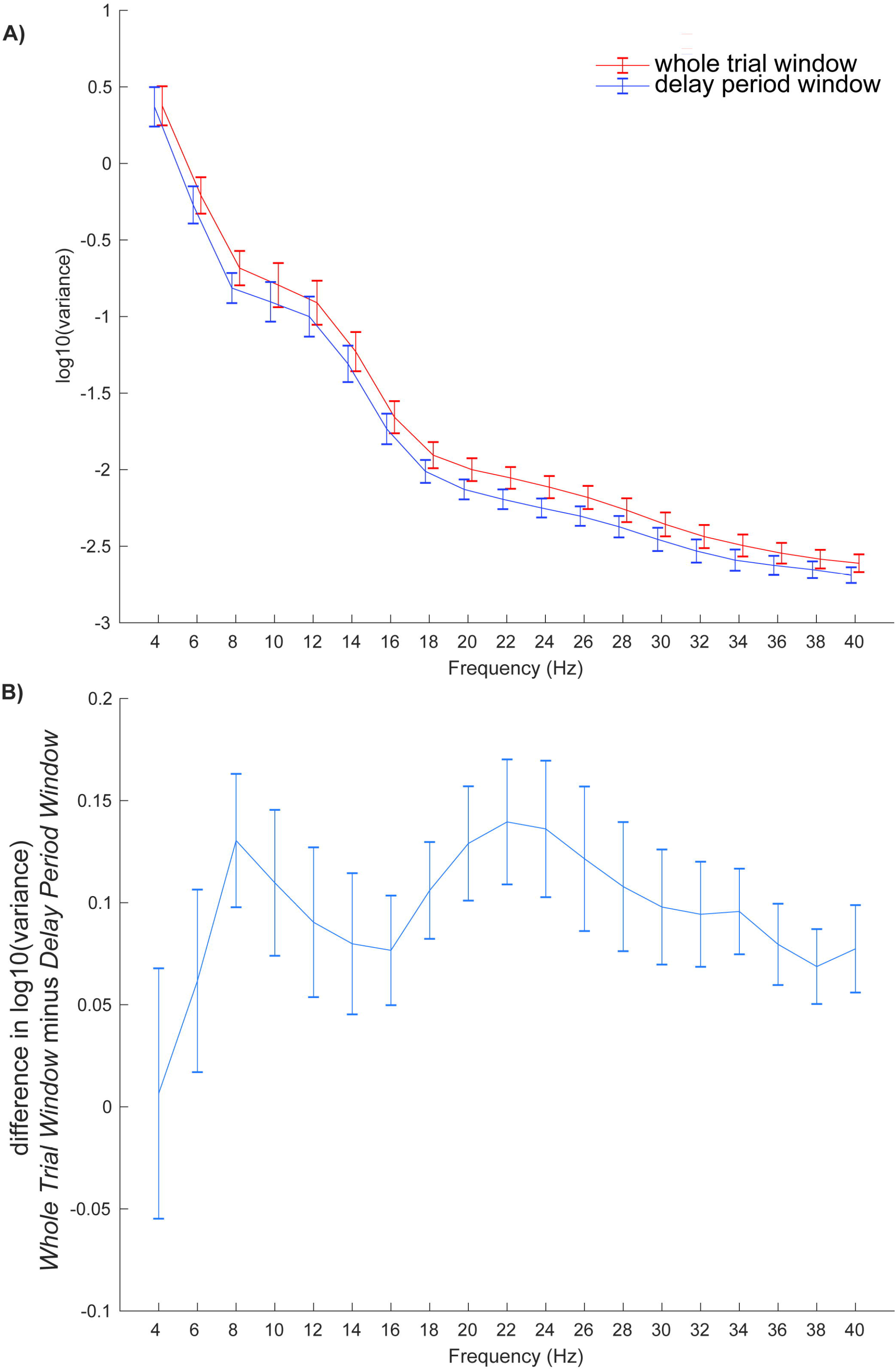
Variance as a Function of Analysis Window and Frequency Bin. The log10 raw amplitude variance (normalized by the N-1 samples) was computed for each subject within two windows - the whole 17 sec trial window (red line) and the 6 sec delay period (blue line) – and plotted as a function of frequency (a). Subject differences in log10 raw amplitude variance (whole trial window minus delay period window) are plotted as a function of frequency (b) and show that whole trial window variances are larger than delay period variances in the range between 6 and 40 Hz, with local peak differences occurring at 8 Hz and 22 Hz. Error bars are subject standard errors of the mean.

### Scrambled Period TSE-Performance Correlations

A hypothesis for why correlations in the delay period disappear when the scrambled scene period is used as a baseline (Fig 7c) is that amplitudes in this period are similar to those found during the delay period, and statistical power is reduced by expression of change relative to a similarly elevated baseline. To explore this hypothesis, a TSE analysis was conducted in sensor space on the 5 sec period of scrambled scene presentation that occurred after the probe choice. There was a cluster of positive correlation found between TSE amplitude and d-prime (red, *Cluster Value* = 660.673, p = 0.017) between 900 ms and 1600 ms after the onset of the scrambled scene (Fig 10a). Individual subject amplitudes expressed as change relative to the mean activity across the 5 sec scrambled period increased as a function of d-prime (Fig 10b), but the range of amplitudes (-0.04 to 0.08) was smaller than the range found for Cluster 2 (-0.01 to 0.24, Fig 6b).

**Figure 10.**
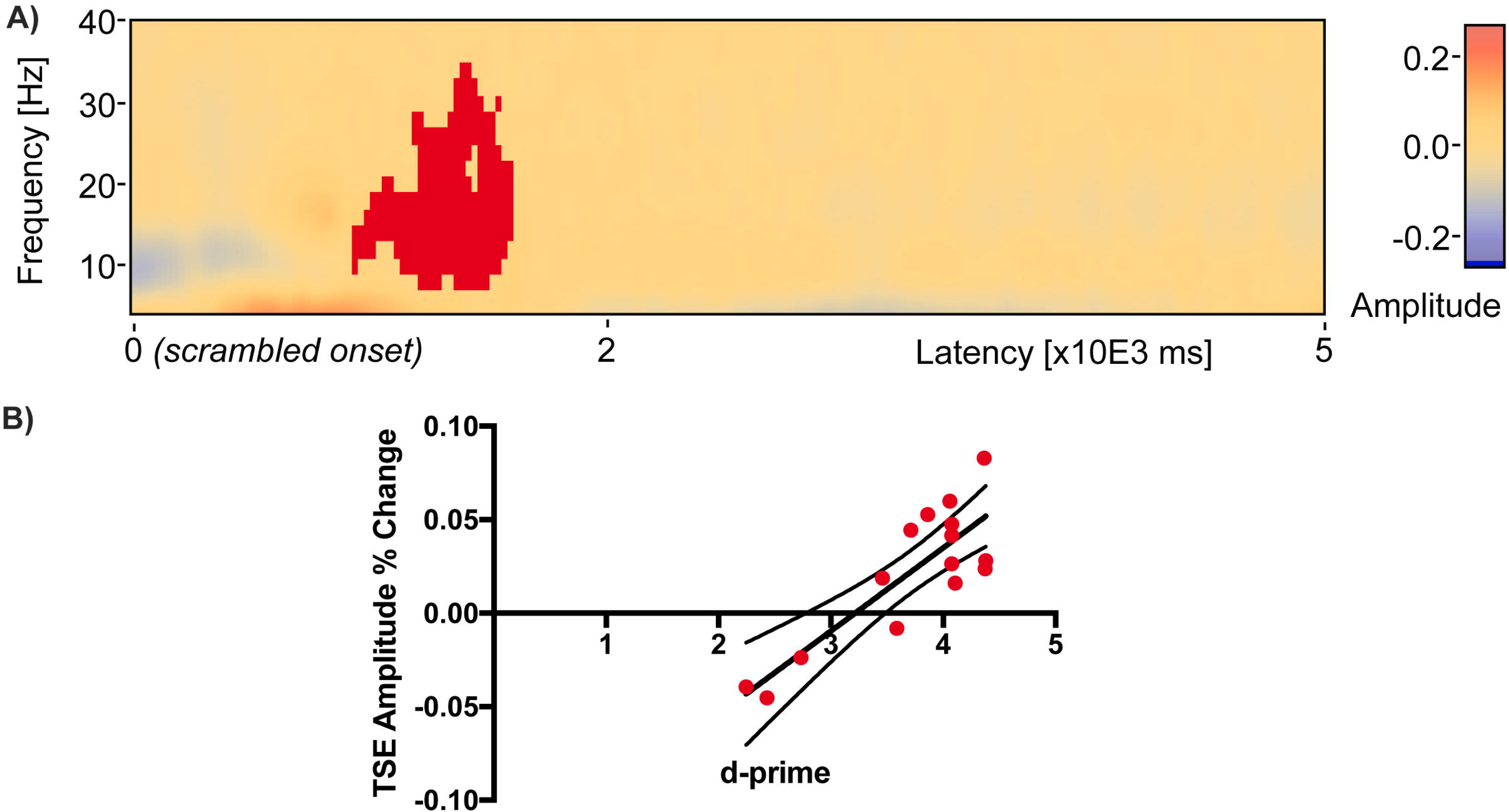
A Significant Cluster of TSE-Performance Correlation in the Scrambled Period. During the 5 second scrambled scene period after probe presentation there was a cluster of positive correlation found between TSE amplitude and d-prime (red, *Cluster Value* = 660.673, p = 0.017) between 900 ms and 1600 ms after the onset of the scrambled scene (a). Individual subject amplitudes expressed as change relative to the mean across the 5 sec scrambled period are plotted as red filled dots with dark line indicating linear regression fits and boundary lines indicating 95% confidence intervals (b).

## Discussion

There are three main findings of the present study. The first is that evidence from the temporal-spectral evolution method suggests transient rather than sustained activity during a relatively long 6 seconds delay period. It has been argued that findings of sustained activity may be a byproduct of averaging over trials and recording locations [7]. The present study, using much longer memory delays, revealed transient patterns even in the trial-averaged data. When examined at both the sensor and source levels, and when using different baselines for comparison, the stable finding that emerged across analyses is an increase in TSE amplitude during the first half of the delay period.

The second main finding is that memory performance is predicted by the early pattern of delay period TSE activity. Early in the delay period, TSE amplitude in the 4-24 Hz range *positively* correlated with d-prime, or the ability of subjects to distinguish old from new scenes. This means that greater TSE amplitude earlier in the delay corresponds with subjects’ ability to distinguish whether a probe scene was one of two scenes presented during the encoding period. Later in the delay period, however, TSE amplitude in the 4-30 Hz range did not correlate positively with performance. In one analysis where the trial window was restricted to the delay period and where the baseline included the same delay period duration, activity *negatively* correlated with d-prime in the latter half of the delay. This pattern of negative correlation toward the latter half of the delay is likely a byproduct of baseline normalization. The early positive cluster, which correlates positively with performance will also correlate with the frequency mean of that trial. Activity toward the end of the delay, which is anti-correlated with behavior, is then normalized by this mean producing a pattern of negative correlation in the latter half of the delay.

The third main finding of the present study is that the patterns of correlation between TSE amplitude and memory performance change when both the temporal window and baseline used for computation of TSE are varied. As noted above, when the temporal window is restricted to just the 6 seconds delay period and the baseline consists of the mean activity across that 6 seconds window, an early positive cluster of correlation is found in addition to two later clusters of negative correlation (Fig 6). The negative clusters are likely a byproduct of TSE normalization using the entire delay period baseline window. When both the window of analysis and baseline are expanded to include the entire trial (Fig 7a), only the single early cluster of positive correlation remains. When using a whole trial window, and restricting the baseline for TSE computation to the 4 seconds encoding period, a similar pattern also emerges (Fig 7b): a single cluster of positive correlation between TSE and performance remains early in the delay period. The decrease in the number of significant clusters of correlations could be caused by decreased statistical power resulting from increased variance as the window of analysis is expanded. A separate analysis of amplitude variance as a function of both the analysis window length and frequency bin revealed increased variance from 6 to 40 Hz in the whole trial window compared to the delay period window. Finally, when using the whole trial window and restricting the baseline for TSE computation to the 5 seconds scrambled scene period following the probe, there are no significant clusters of either positive or negative correlation anywhere in delay period or anywhere in the trial window. A separate analysis of activity in the 5 sec scrambled period window revealed a significant cluster of elevated amplitude change that was positively correlated with performance, similar to the cluster of positive correlation early in the delay period but corresponding to a reduced range of amplitudes. This shows that TSE amplitudes during the period of scrambled scene presentation following the probe choice vary similarly to activity during the early portion of the delay period (Fig 10b versus Fig 6c, Cluster 2).

The increase in TSE amplitude early in the delay period is consistent with event-related synchronization, with the event or time-locking being the start of the delay period [36, 42]. One interpretation of increased synchronization early in the delay period is that it reflects a turning of attention inward after the visual input stops. This pattern is consistent with cognitive processing associated with the attempt to maintain the two previously presented scenes in an online buffer. The speculated attempt to hold the scenes in an online buffer, even if maintenance only lasted for a few seconds [43, 44] may have helped performance, as behavior correlated positively with activity in this period. Maintenance for a short period of time followed by decay is consistent with the observed elevated activity early in the delay, only lasting for the first 2 or 3 seconds followed by the tapering of activity [4, 15, 16, 19, 20, 45]. This interpretation is supported by the finding of a positive correlation of only the early TSE amplitude with subsequent memory performance: the higher the TSE early in the delay period, the better subjects are at distinguishing old from new scenes during the subsequent probe phase. Alternatively, activity early in the delay might have been related to encoding into episodic memory and therefore positively correlated with behavior. If this type of encoding is successful, working memory would not have to be maintained which could explain the non-stationary delay activity.

The decrease in TSE amplitude toward the end of the delay period is consistent with event-related desynchronization, with time-locking being the beginning of the delay period [36, 42]. One interpretation of this late pattern of activity is that it reflects the motor planning that subjects make in anticipation of the end of the delay [37, 46]. The greater the decrease in TSE amplitude during the end of the delay period, the better subjects are at making a button press after distinguishing old from new scenes during the probe phase. An alternative to the motor planning interpretation is that this desynchronization instead reflects a reactivation of the encoded scenes late in the delay period as subjects anticipate the approaching probe comparison [47, 48]. This alternative reactivation interpretation is supported by a similar pattern of negative correlation between TSE amplitude and memory performance during the actual encoding period, when the stimuli are visibly present, that was obtained in the source analysis in the right parietal region (Fig 8b).

An additional novel contribution of this report is the demonstration that changing the temporal window of analysis and baseline can alter the TSE-performance correlations obtained. The most extreme example in the sensor analyses performed here occurred when the baseline for the TSE analysis was switched to the 5 seconds period of scrambled scene presentation occurring after the probe scene was presented (Fig 7c). All clusters of significant correlation between TSE and memory performance disappeared, even though the pattern of higher TSE early in the delay and lower TSE during encoding and around the probe remained. One explanation for this is that, as in the early portion of the delay period, during the presentation of scrambled scenes subjects’ attention is turned inward and focused on the previously presented scenes. Thus, when the scrambled period is used as a baseline, it effectively cancels out the similar activity during the delay period. During the viewing of scrambled scenes immediately after the probe, it is possible that the EEG signal reflects patterns of activity similar to the activity present early in the delay period. This is plausible especially for the 50% of the trials where the immediately preceding probe was a positive probe, and could have reminded the subject of one of the two previously encoded scenes. This scenario is one explanation for why using the scrambled period of activity as a baseline reduced the TSE delay activity correlation with memory performance.

This transient pattern of activity that was observed during the delay period is inconsistent with some previous research [4, 15, 16, 19, 20, 45]. Studies using MEG [49–51] and EEG [52, 53] have found sustained, not transient, increases in alpha power (8-12 Hz) during the delay period for stimuli that were successfully remembered compared to forgotten stimuli. These studies have also reported sustained activity in the theta range (4-8 Hz) [19, 20, 45]. Increases in beta oscillations (12-30 Hz) have also been observed throughout the delay period of working memory tasks [4, 51]. In addition, sustained decreases in alpha and beta power are typically associated with semantic encoding of stimuli [54] and similar decreases in theta power have also been observed throughout task encoding [55] but not retention periods.

More recently, delay activity has also been associated with enhanced activity in the medial temporal lobe (MTL) [56]. According to one framework [47], synchronization and desynchronization reflect a division of labor between hippocampus, a structure in MTL, and neocortex respectively. In the neocortex, information is stored in material-specific regions, while the hippocampus binds information among widespread cortical regions. In this framework, it is postulated that synchronization and desynchronization mechanisms interact in the formation of episodic memories, but the same processes may be at work during the initial processing of novel stimuli in working memory. Information transmission between neocortex and hippocampus is believed to be related to activity in the theta range, and more specifically the cross coupling of theta and gamma oscillations [47, 57, 58]. Within one theta cycle, different patterns of neurons activate during each gamma cycle, and represent an object that is held in memory. Multiple objects can be held together, each represented by a different gamma cycle [59, 60]. Moreover, phase synchronization of theta activity between prefrontal and posterior brain regions reflects engagement of attention or central executive systems [56]. Thus, the observed theta activity may also reflect communication between the medial temporal structures and neocortex to facilitate the binding of encoded information for later retrieval. Alpha and beta activity observed during working memory processes are also implicated in the processing and storage of information. Decreased activity in both the alpha and beta range is related to the indexing of information in cortex during information processing. Reactivation is associated with activity in the same material-specific regions where activity was observed during initial processing [61]. Because desynchronous alpha and beta activity appear to be one correlate of a mechanism for storing information during memory encoding, observation of this activity during the later portion of the delay period raises the question of whether it may represent reactivation of encoded stimuli in preparation for comparison with a subsequent probe stimulus.

In the present study, the correlation of delay activity with memory performance was affected by changing the baseline period used in the TSE computation. Patterns of significant correlations were observed most clearly when the entire delay period itself served as its own baseline. When the scrambled scene period was used as the baseline, the significant activity correlations with performance disappeared. The scramble scene period after the probe is a time when the participant may be directing attention inward and mostly ignoring visual input [42]; hence, it could be argued that this baseline resembles a pre-stimulus baseline in anticipation of another upcoming set of stimuli to be encoded. Alternatively, the scrambled scene occurred immediately after the probe, a time during which on at least half of the trials the subject would have been just reminded of one of the encoded stimuli. Therefore, reactivation of the encoded stimuli could be reflected in the activity during this scrambled scene period, and this activity could be similar to the patterns early in the delay period. Such a scenario would explain why delay activity performance correlations disappeared when the scrambled scene is used as a baseline. Many previous studies employ a baseline period that occurs before the presentation of stimuli [35, 36] and participants are instructed to focus on a fixation cross [4, 49, 62]. Recently it has been suggested that a baseline period in which the participant is not engaged in any cognitive activity could bias the interpretation of the spectral changes [63]. Therefore, future studies should examine how cognitive processes interact with different visual inputs in baseline periods before encoding, during the delay period, and after probe presentation.

Previous studies have often utilized relatively short retention periods of approximately 3 seconds or less [4, 15, 16]. The present study used a delay period twice as long. Yet, the pattern of synchronous activity that was found early in this long delay period had a duration (~3 seconds) that was similar to the aforementioned studies. Extending the delay period in the present study resulted in activity that is temporally restricted to only a few seconds, but about the same length as reported in other studies. One interpretation is that the delay period durations used in previous EEG studies may have been too short to reveal transient, early activity. In addition, many studies that examine delay activity have excluded activity from trials where the subject subsequently made incorrect responses. In this study, both correct and incorrect trials were included in the TSE matrix computations to be able to compute accurately correlations with the d-prime performance sensitivity measure, which is computed based on both correct and incorrect trials. It would not be optimal to correlate only correct trial activity patterns with a measure of performance computed from both correct and incorrect behavioral choices. It has been previously reported that correct and incorrect responses produce different patterns of delay activity [64]. The likely explanation for the correlation between performance and activity across subjects reported here is that subjects had differences in activity and strategies for task performance that lead to a difference in their performance, rather than the finding being driven directly by different activity during correct and incorrect trials.

Finally, the present study utilized complex naturalistic stimuli, as compared to many studies that have employed simple shapes, letters, or numbers. Some working memory studies have used faces [16, 49] or real-world scenes [65]. Although the early pattern of activity exhibited during the delay is consistent with studies that have used simple stimuli, it was not sustained throughout the entire delay period. The early positive activity that faded after 3 seconds may reflect visual memory decay associated with the difficulty of maintaining complex scenes without effortful covert rehearsal. If subjects had difficulty attaching a verbal label to the complex stimuli, they may not have been able to employ rehearsal mechanisms. While some stimuli were easier to assign a verbal label (e.g., scene with a building), other stimuli may have been more difficult (e.g., scene with trees, a valley, and mountains). When memory for stimuli that were difficult to label began to fade, underlying synchronous activity may have become more desynchronous around 3 seconds. If on the other hand, participants were able to verbally label the scenes and rely on rehearsal, they may have been able to maintain the stimuli throughout the retention period thus resulting in a sustained pattern of synchronous activity through the whole delay period. It is worth noting that in the source analyses, there were no significant correlations of TSE delay activity with performance in any left frontal region, the brain area implicated in covert articulatory rehearsal. Rather, the only significant TSE activity performance correlation was obtained in the right parietal region. This region has is consistent with the neural substrate of the working memory visuospatial sketchpad’s inner scribe and visual cache. The phonological loop’s region of articulatory rehearsal [1] is typically localized to left frontal regions. Future source estimation studies should examine the transient versus sustained nature of the delay activity when the phonological loop is recruited during covert rehearsal with explicit verbal labels and also during control conditions with and without articulatory suppression (e.g., repeating ‘the’ instead of a scene-based verbal label).

## Limitations

All measurement techniques and analysis methods have limitations, and the present study is no exception. The TSE technique involves analysis of signals around an event of interest, which in this study, was the onset of the delay period. One could argue that synchronization is more likely to manifest closer to delay onset, with desynchronization more likely as a function of increasing time from the delay onset event. This could explain the average sensor space finding of more synchronized positive amplitudes near delay onset with more desynchronized negative amplitudes after 3 sec of maintenance. We note, however, that an exception to this pattern is a very early burst of desynchronization in the first 500 ms between 4 and 18 Hz (Fig 6a, blue). Regarding the extent of clusters in time, it should be taken into consideration that the duration of an event-related potential or field is dependent on other factors beside the underlying neural activity in a cortical region. That means a brain response can be more or less time-locked to a stimulus (or delay) onset, leading to very sharp (highly time-locked) or more wide-spread (more jittered) potential responses or fields. The delay period sensor space analysis, which involved averaging over all electrodes and trials, allowed for the most statistical power. When the analysis window was expanded, statistical power was likely reduced as analysis of raw amplitudes showed increased variance as a function of window length (Fig 9). While EEG allows for millisecond temporal resolution, attenuation of the signal through meninges, skull and scalp reduces signal to noise and limits spatial resolution. There were no MRI data available for the subjects studied here so only the most basic source estimation was performed, which also reduced statistical power relative to the sensor space analysis by dividing source space into fifteen different brain partitions.

Finally, there is a limitation in terms of what one can infer about the type of cognitive processing subjects used in the present study. Humans are adept at encoding scenes and can remember many scenes from minutes to days after initial presentation. We did not design this study to test a cognitive hypothesis for scene processing and therefore neither encouraged or dissuaded subjects from using a purely verbal or purely visual rehearsal strategy. Based only on informal debriefings from some subjects can we speculate that there is variability about how easy it is to apply verbal labels to these scenes. Similar to the literature on mental imagery, some subjects might be more adept at quickly assigning labels in the two seconds they have to view each scene, while others would likely find such a task more challenging and effortful. Unfortunately, we do not have data on which scenes are easy to label and which are not. Such ratings are not part of the SUN database and we did not gather any subject ratings indicating the ease with which they could apply labels to these different stimuli. Nevertheless, future studies should investigate the ease of assigning a verbal label as a maintenance strategy and the impact of using such a strategy on delay activity. Since the stimuli were complex color scenes, which were difficult to label and therefore not amenable to verbal rehearsal, one strategy subjects might use to correctly recognize the scenes is familiarity detection. A strategy of familiarity is a possibility, especially for the relatively long delay of 6 sec when maintenance using a verbal strategy is not straightforward. Rather than seeing temporal activity in the source analyses, which would be more consistent with a familiarity strategy, or prefrontal activity, which would be consistent with central executive working memory, we found parieto-occipital activity. More controlled behavioral studies will need to be conducted to determine whether or not this spatial pattern of activity reflects a strategy of short-term storage in a visuospatial sketchpad consistent with Baddeley’s multi-compartment model or a strategy of familiarity detection. Finally, we did not vary delay period duration within or across subjects. Had delay duration been varied from 2 to 6 sec it is possible stationary activity would have been obtained, with non-stationary activity manifesting as delay length increased to 6 sec. A potential explanation for less sustained activity in the present study relative to other studies might be the relatively longer delay. Future studies should vary delay period duration within subjects to determine if non-stationary activity is revealed at long delays, and stationary activity obtained at short delays.

## Conclusions

Analysis of the temporal spectral evolution of the delay period EEG suggests that short-term visual memory maintenance involves a period of transient activity in which amplitude increases in the first half of a 6 second fixed delay period contribute to subsequent task performance. Early in the delay period these TSE amplitude changes are positively correlated with subsequent successful short-term memory for scenes. The significance of the early delay activity-performance correlations depends on the trial time window and baseline periods used in the analysis. These results add to the growing literature supporting transient rather than sustained activity changes during working memory delay periods. Additional research is needed to understand how different cognitive processes involved in maintaining complex visual information modulates underlying oscillatory activity, and how synchronous and desynchronous patterns reflect intra - and inter-regional information transmission.

## Acknowledgements

The authors thank Miriam San Lucas, Ning Mei, and Ben Fernandez for assistance with data collection and analysis.

## Author Contributions

Conceived and designed the experiments: T.M.E.

Collected data: C.R. and K.N.

Analyzed data: C.R. and K.N.

Wrote the paper: T.M.E. and C.R.

